# Neurons in the lateral spinal nucleus give rise to the spinotectal component of the anterolateral system in the mouse

**DOI:** 10.64898/2026.09.29.755366

**Authors:** Mai Abu Hajer, Mansi Yadav, Wenhui Ma, Allen C. Dickie, Erika Polgár, Andrew M. Bell, Andrew J Todd

## Abstract

The anterolateral system (ALS) originates from neurons in various regions of the spinal cord and conveys somatosensory information related to pain, skin temperature and itch to several brain targets. One component of the ALS is the spinotectal tract, which projects to the superior colliculus (SC) and is thought to underlie directed movements of the head towards a painful stimulus. Using anterograde and retrograde tracing, we demonstrate that the input from the spinal enlargements to the SC originates predominantly from neurons in the lateral spinal nucleus (LSN), many of which express calbindin. This input, which mainly targets the caudal and lateral parts of the SC, is somatotopically organised, and is associated with the clusters of Pitx2-expressing neurons that have been identified in previous studies. Since activation of Pitx2 neurons drives directed head movements, input from LSN neurons to the Pitx2 cells is likely to be a critical component of orientation and attention towards a noxious stimuli. We also provide evidence that spinotectal LSN cells receive excitatory synaptic input from neurons in superficial dorsal horn that express substance P or the gastrin-releasing peptide receptor (GRPR). Since GRPR neurons are strongly implicated in itch, our findings raise the possibility that the SC has a role in coordinating movements in response to pruritic stimuli.

## INTRODUCTION

The superior colliculus (SC), located in the dorsal part (tectum) of the midbrain, registers events in surrounding space through visual, auditory and somatosensory channels and is involved in both orientation and avoidance movements in response to external stimuli^1–4^. The SC is multilayered, and receives sensory information in a highly organised way. Visual inputs target the superficial layers, while auditory and somatosensory inputs innervate the intermediate and deep layers. SC neurons with somatosensory responses can be activated by low-threshold (touch and hair movement) and/or noxious stimuli^5–9^, indicating that the SC receives input originating from both low-threshold mechanoreceptors and nociceptors. The low-threshold input travels via the gracile and cuneate nuclei, which are innervated by Aβ cutaneous low-threshold mechanoreceptors^10^ and send axon collaterals the SC^11^. In contrast, nociceptive input is thought to be mainly from spinal projection neurons belonging to the anterolateral system (ALS)^12–16^.

The ALS underlies perception of pain, itch and skin temperature, as demonstrated by the loss of these sensations following anterolateral cordotomy^17–19^. ALS neurons project to multiple brain regions, including the reticular formation, the nucleus of the solitary tract (NTS), the lateral parabrachial area (LPB), the periaqueductal grey matter (PAG), the SC and various thalamic nuclei^14,15,20–23^, with individual ALS neurons innervating several different brain areas^13,24^. ALS projection neurons are unevenly distributed within the spinal cord, being concentrated in lamina I, the lateral spinal nucleus (LSN), the lateral part of lamina V and the region around the central canal^10,12,15,25–27^. Emerging evidence indicates that different types of ALS cells have characteristic patterns of projection to the brain^13,20,21,23,28^. For example, somatostatin-expressing ALS neurons in lateral lamina V^29^ project directly or indirectly to the mediodorsal nucleus of the thalamus^23^, suggesting a role in the motivational/affective dimension of pain^30^, while cold-selective ALS neurons in lamina I project to brainstem and thalamic regions involved in cold perception and thermoregulation^28^.

The projection from ALS neurons to the SC (the spinotectal tract) is thought to underlie orientation of the head towards a painful stimulus, for example to allow licking of the injured region^8,13,31–33^, an integral feature of the response of many mammals to acute or inflammatory pain^34–36^. Despite its importance, little is known about the cells of origin of the spinotectal tract.

Early retrograde tracing studies investigated projections from spinal cord to the midbrain, but in some cases injection sites included the PAG^37–42^, which receives a substantial projection from spinal cord. In a recent study, we found that intraspinal injections of AAV coding for Cre-dependent tdTomato in *Calb1*^Cre^ mice gave axonal labelling in the SC, but only when the injections included the LSN^28^. Since calbindin, which is encoded by *Calb1*, is expressed by many ALS neurons in the LSN^28,43^, this suggested that projection neurons in the LSN may be an important source of the spinotectal tract. Here we have tested this hypothesis, looked for evidence that would support a role for the spinotectal tract in directed movements of the head towards a noxious stimulus, and investigated sources of excitatory synaptic input to spinotectal neurons in the LSN.

## RESULTS

### Anterograde viral labelling in the *Calb1*^Cre^ mouse

As part of a previous study^28^ we had injected AAV1.CAG.Cre^ON^.tdTomato into the L3-L5 segments of 3 *Calb1*^Cre^ mice. When injection sites included the LSN (see Figure 6-figure supplement 1 from reference^28^), we observed anterograde axonal labelling in the contralateral SC. Here we explore this labelling in more detail. As shown in Figure 1, the labelled axons occupied the caudal half of the SC, extending from ∼4.8 to 4.0 mm caudal to the bregma^44^. They were located within the intermediate grey matter, and labelling was densest at the extreme caudal end of the SC, where it formed a continuous band that occupied the lateral part of the colliculus (Figure 1D,E).

**Fig 1.**
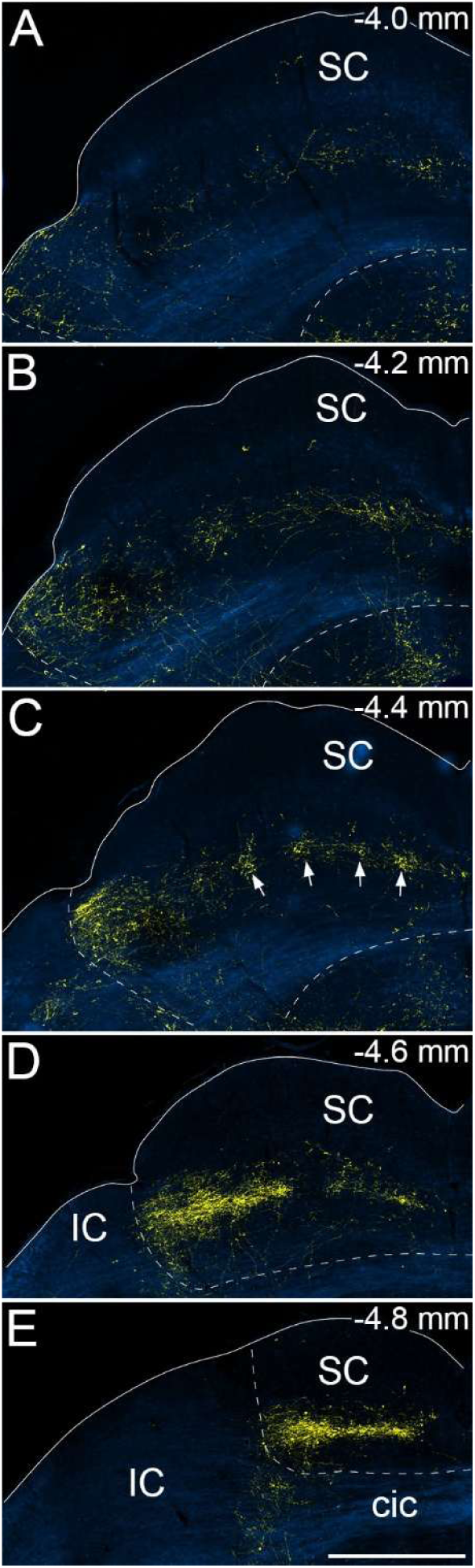
Axonal projections from calbindin-positive neurons in the lumbar enlargement to the superior colliculus (SC). The images show labelling with tdTomato (yellow) superimposed on dark field images (blue) at progressively more caudal levels (**A**-**E**) of the contralateral superior colliculus from a *Calb1*^Cre^ mouse that had received intraspinal injections of AAV1.CAG.Cre^ON^.tdTomato into the L3, L4 and L5 segments on one side. Estimated levels are indicated as millimetres caudal to the bregma and approximate borders of the SC are shown with dashed lines. There is dense labelling in the most caudal sections (**D**,**E**), particularly in the lateral part of the SC, and this diminishes at more rostral levels (**A**-**C**). In some sections there are clear clusters of axonal labelling, and this is well seen in **c**, where the clusters are marked by arrows. A similar pattern of labelling was seen in 2 other *Calb1*^Cre^ mice that had received intraspinal injections of AAV1.CAG.Cre^ON^.tdTomato into the same segments. cic, commissure of the inferior colliculus; IC, inferior colliculus. Scale bar = 500 µm.

Labelling became progressively sparser at more rostral levels, and in some sections it could be seen to form discrete patches within the intermediate zone (Figure 1B,C) as noted in previous studies^14,15^.

Anterogradely labelled axons were also seen in the corresponding regions of the ipsilateral SC at a lower density.

Masullo et al^45^ identified a population of neurons in the intermediate grey layer (stratum griseum intermediale, SGI) that expressed the transcription factor Pitx2, and showed that activating these cells drove 3-dimensional head displacements. The Pitx2 cells were clustered into “modules”, forming a honeycomb arrangement that spanned the width of SGI, with somata located in the ventral part of this layer and dendrites that occupied its full thickness. These modules appeared to have a similar distribution to the patches of axonal labelling observed here and in previous studies^14,15^. To compare the distribution of spinal input with that of Pitx2-expressing neurons, we therefore carried out fluorescence *in situ* hybridisation on sections from 3 *Calb1*^Cre^ mice that had received intraspinal injections of AAV1.CAG.Cre^ON^.tdTomato into the lumbar spinal cord. We found that the patches of axonal input were often aligned with clusters of Pitx2 neurons, although the axons generally lay dorsal to the Pitx2 cell bodies (Figure 2).

**Fig 2.**
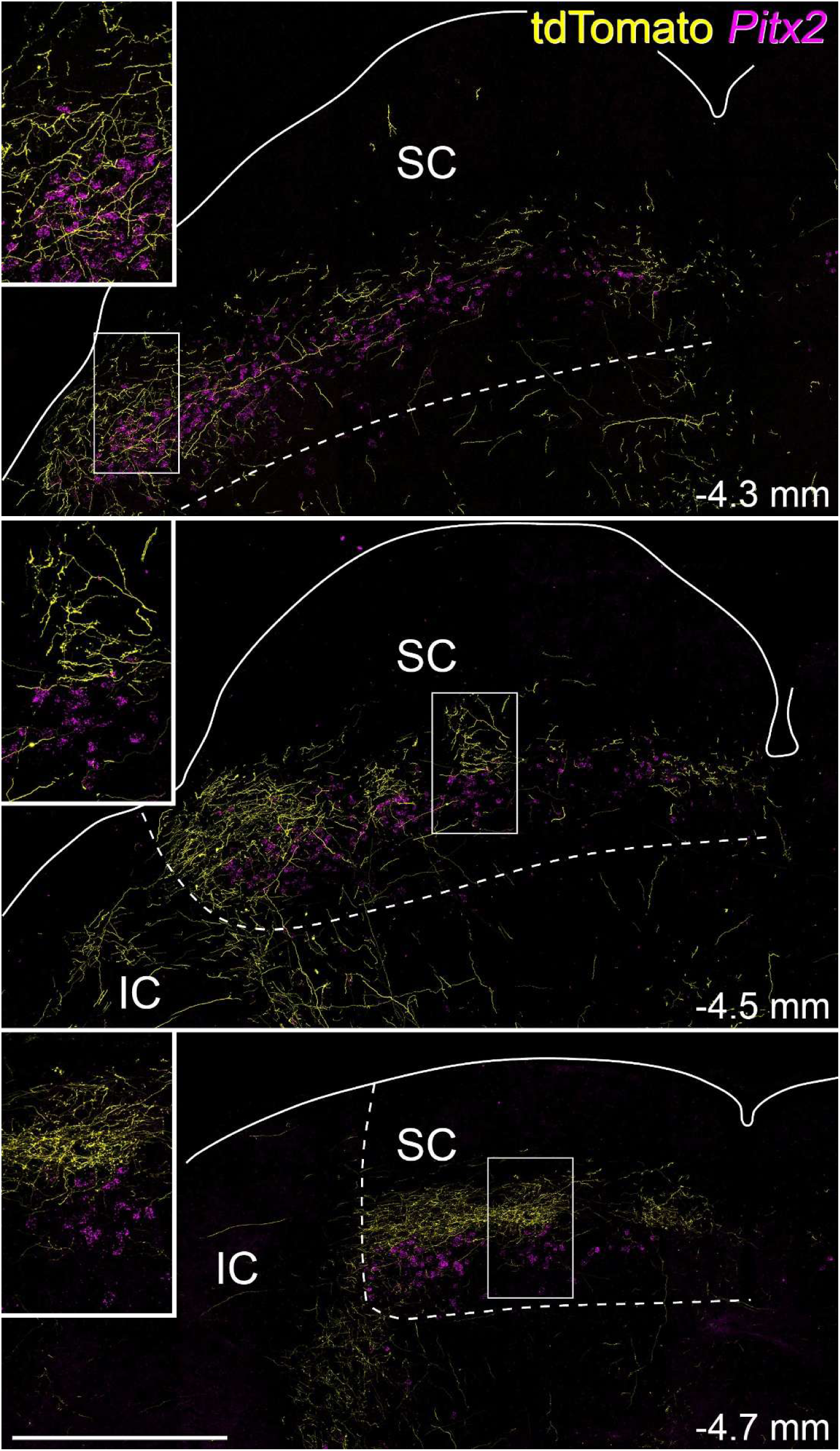
Relationship between clusters of spinotectal axons and Pitx2 expression in the superior colliculus from 3 *Calb1*^Cre^ mice injected with AAV1.CAG.Cre^ON^.tdTomato in the L3, L4 and L5 spinal segments. The sections have been processed for fluorescence *in situ* hybridisation to reveal *Pitx2* mRNA (magenta) and immunohistochemistry to show tdTomato-labelled axons (yellow). The dashed lines show the approximate borders of the superior colliuculus (SC). Each section is from a different mouse, and at slightly different levels through the colliculus, with estimated distances caudal to Bregma indicated in each case. As shown in Figure 1, axonal labelling is densest in the lateral part of the superior colliculus and forms clusters more medially. *Pitx2*-positive cell bodies are generally located slightly ventral to the labelled axons, and in some regions clusters of axons and clusters of *Pitx2* cells are aligned. Examples of this are shown in the insets (corresponding to the boxed region in each image). Scale bar = 500 µm.

To look for evidence of a somatotopic pattern of labelling, we initially examined SC inputs that resulted from injection of AAV1.CAG.Cre^ON^.tdTomato into the C8 segment of 3 *Calb1*^Cre^ mice, again in tissue that was generated as part of a previous study (see Figure 6-figure supplement 3 from reference^28^). The axonal labelling in SC was sparser than that seen following injections into the L3-5 segments (presumably at least partly due to the more restricted spinal injection) and appeared to be displaced rostrally, extending from ∼4.8 to 4.0 mm caudal to bregma, but with the maximum density at 4.4-4.2 mm caudal to bregma. As for lumbar injections, it was strongest in the lateralmost part of the SC at these levels and patchy labelling was seen in some sections (Figure S1).

To provide more direct evidence of a somatotopic organisation, we made injections of AAV9.CB7.GFP into the L3 segment and AAV9.CB7.mCherry into the C8 segment in 4 wild-type mice. Axonal labelling in the SC was stronger than that seen after injection of AAV1.CAG.Cre^ON^.tdTomato in *Calb1*^Cre^ mice, and this may reflect stronger fluorescent protein expression from the viruses used for this part of the study, as well as the fact that not all spinotectal neurons express calbindin (see below). Consistent with the findings from the single injections, we found that the GFP labelling (from the lumbar injection) was strongest at the extreme caudal end of the SC at levels between 4.8 and 4.6 mm caudal to bregma, while mCherry labelling (from the cervical injection) became prominent at more rostral levels – 4.4 to 4.0 mm caudal to bregma (Figure 3). Again, the labelling was strongest in the most lateral parts of the SC. Despite the differences in rostrocaudal extent of anterograde labelling from the two injection sites, there was considerable overlap (Figure 3). Interestingly, in sections with patches of labelled axons these frequently contained both mCherry- and GFP-positive axons, indicating input from both cervical and lumbar enlargements to the same patch (Figure S2).

**Fig 3.**
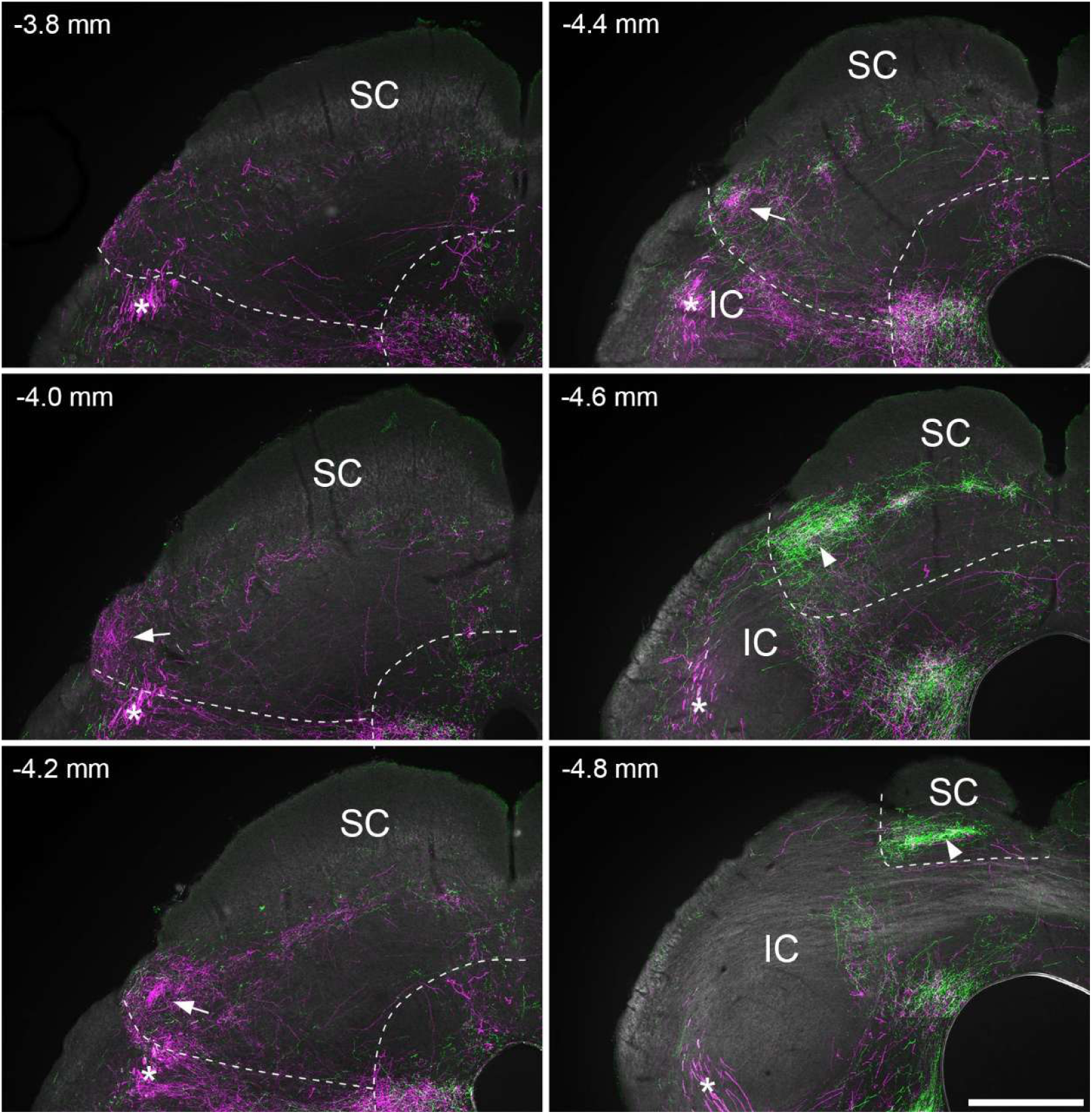
Axonal projections from lumbar and cervical enlargement to the superior colliculus. The images show labelling with GFP (green) and tdTomato (magenta) in the left (contralateral) superior colliculus (SC) following injections of AAV.CB7.GFP into the L3 segment and AAV.CB7.mCherry into the C8 segment on the right side. Estimated levels are indicated in millimetres caudal to the bregma and the approximate borders of the SC are shown with dashed lines. There is strong GFP labelling in the caudal-most sections (-4.8 and -4.6), located in the lateral part of the SC (arrowheads). mCherry labelling is present in more rostral sections (between -4.0 and -4.4), and again, this is densest in the lateral part of the SC (arrows). Patches of GFP and mCherry axonal labelling can be seen in the central part of the SC at -4.4, and these are shown in more detail in Figure S2. A prominent bundle of mCherry-labelled axons (marked with asterisks) can be seen just outside the SC, and since this could be followed rostrally into the thalamus, it presumably corresponds to the spinothalamic tract. A similar pattern of labelling was seen in 3 other wild-type mice that had received equivalent intraspinal injections of AAV.CB7.GFP into L3 and AAV.CB7.mCherry into C8. IC, inferior colliculus. Scale bar = 500 µm.

### Retrograde labelling following CTB injections into the SC

To test the prediction that spinal projections to the SC originated mainly from the LSN, we carried out retrograde labelling experiments in which cholera toxin B subunit (CTB) was targeted to the SC. We injected small amounts (100 nl of 1% CTB) in 4 wild-type mice, resulting in injection sites that were restricted to the SC (Figure S3). We examined retrograde labelling in transverse sections through the L4 and C7 segments, representing the lumbar and cervical enlargements, as well as in transverse sections of the C2 segment. C2 was chosen for two reasons, firstly because it contains the lateral cervical nucleus (LCN), which also projects to the SC^40,46,47^, and secondly to allow comparison with data from a recent study in which fluorescence micro-optical section tomography (fMOST) was used to reconstruct the axons of projection neurons in the upper cervical (C1-C4) segments of the mouse^13^.

In the L4 segment, we found the highest numbers of retrogradely labelled cells in the LSN, with more cells on the side contralateral to the injection site (7.8 cells/10 sections in contralateral LSN, 3.6 cells/10 sections in ipsilateral LSN) (Table 1, Figures 4, 5). The CTB labelling in individual cells was largely restricted to the cell body, with very limited extension into primary dendrites.

**Fig 4.**
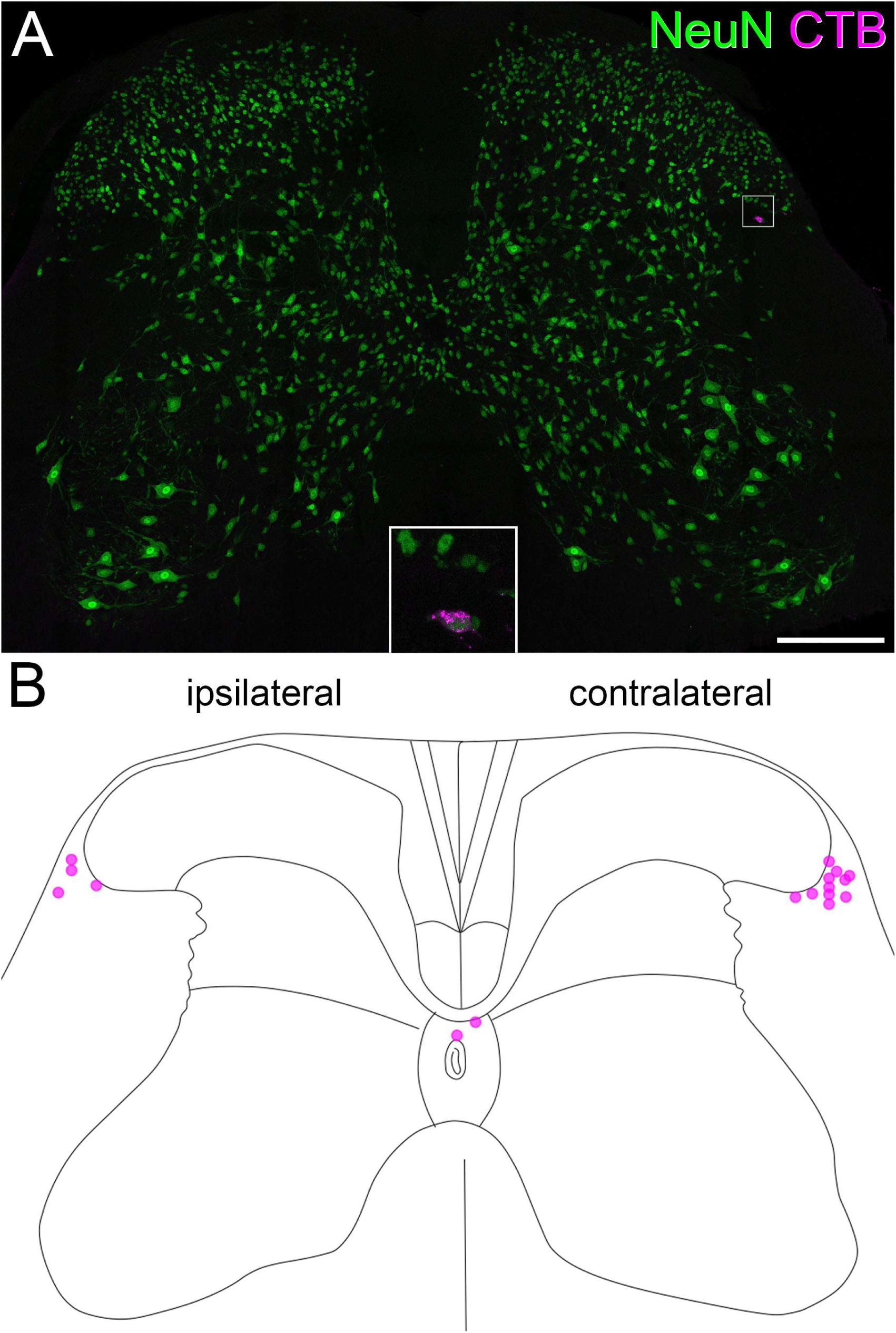
Retrograde labelling in lumbar spinal cord from an injection of CTB that was limited to the superior colliculus. **A**: A section through the L4 segment immunostained to reveal the neuronal marker NeuN (green) and CTB (magenta). A single retrogradely labelled cell is present in this section, and this is located in the LSN on the side contralateral to the superior colliculus injection. The area within the box, which contains the retrogradely labelled cell, is shown at higher magnification in the inset. Note that CTB occupies part of the perikaryal cytoplasm, and that the nucleus of this cell is labelled by the NeuN antibody. **B**: A plot of all retrogradely labelled neurons present in fourteen 60 µm thick transverse sections from this experiment. Ipsilateral is shown to the left and contralateral to the right. Each circle represents a single labelled cell, and the outline was taken from the Allen Brain Atlas (https://mousespinal.brain-map.org/imageseries/showref.html). The CTB injection site for this experiment is shown in Fig S3B. Scale bar = 200 µm.

**Table 1.** Quantitative data from injections of CTB that were restricted to the SC

|  |  | Contralateral |  |  |  |  | Ipsilateral |  |  |  |  |
| --- | --- | --- | --- | --- | --- | --- | --- | --- | --- | --- | --- |
|  | # sections | LSN | LCN | Lamina I | DDH | CC | LSN | LCN | Lamina I | DDH | CC |
| C2 | 13.8<br>(7-19) | 6.28<br>(3.57-8.42) | 6.38<br>(1.33-15.71) | 2.93<br>(0-4.74) | 2.22<br>(0.71-4.29) | 0 | 0.95<br>(0-1.43) | 1.81<br>(1.05-3.33) | 0.13<br>(0-0.53) | 0.85<br>(0-1.43) | 0 |
| C7 | 11.3<br>(9-14) | 6.56<br>(5.56-7.86) |  | 0.89<br>(0-1.43) | 1.14<br>(0-2) | 0 | 3.56<br>(2.5-4.44) |  | 0.43<br>(0-1) | 0.49<br>(0-1.11) | 0 |
| L4 | 14.8<br>(14-17) | 7.74<br>(5.71-11.18) |  | 0.35<br>(0-0.71) | 1.19<br>(0-2.14) | 0.36<br>(0-1.43) | 3.6<br>(0.71-8.46) |  | 0 | 0.15<br>(0-0.59) | 0 |

Column 2 shows the mean numbers of sections analysed for each of the 4 mice with the range in brackets. Columns 3-11 show mean numbers of cells retrogradely labelled with CTB per ten 60 µm transverse sections (range in brackets). CC, area around central canal; DDH, deep dorsal horn; LCN, lateral cervical nucleus; LSN, lateral spinal nucleus.

Although a few retrogradely labelled cells were seen in the deep dorsal horn and the area around the central canal, these were far less numerous (Table 1), and cells in the contralateral or ipsilateral LSN accounted for 85% of all retrogradely labelled neurons (11.34 cells per 10 sections, compared to 2.05 cells per 10 sections for all other regions, Table 1). A similar pattern was seen in the C7 segment, with the highest numbers of cells present in the contralateral LSN, and several in the ipsilateral LSN, as well as a few labelled cells in lamina I and the deep dorsal horn (Figure 5, Table 1). However, as for L4, neurons in the ipsilateral or contralateral LSN accounted for the great majority (77%) of retrogradely labelled cells in the C7 segment (Table 1). Note that these numbers are likely to be an underestimate of the total spinotectal projection as it is not possible to fill the whole of the region of SC that contains spinotectal axons with CTB, without spread of tracer into other areas that also receive input from the spinal cord (e.g., the PAG).

**Fig 5.**
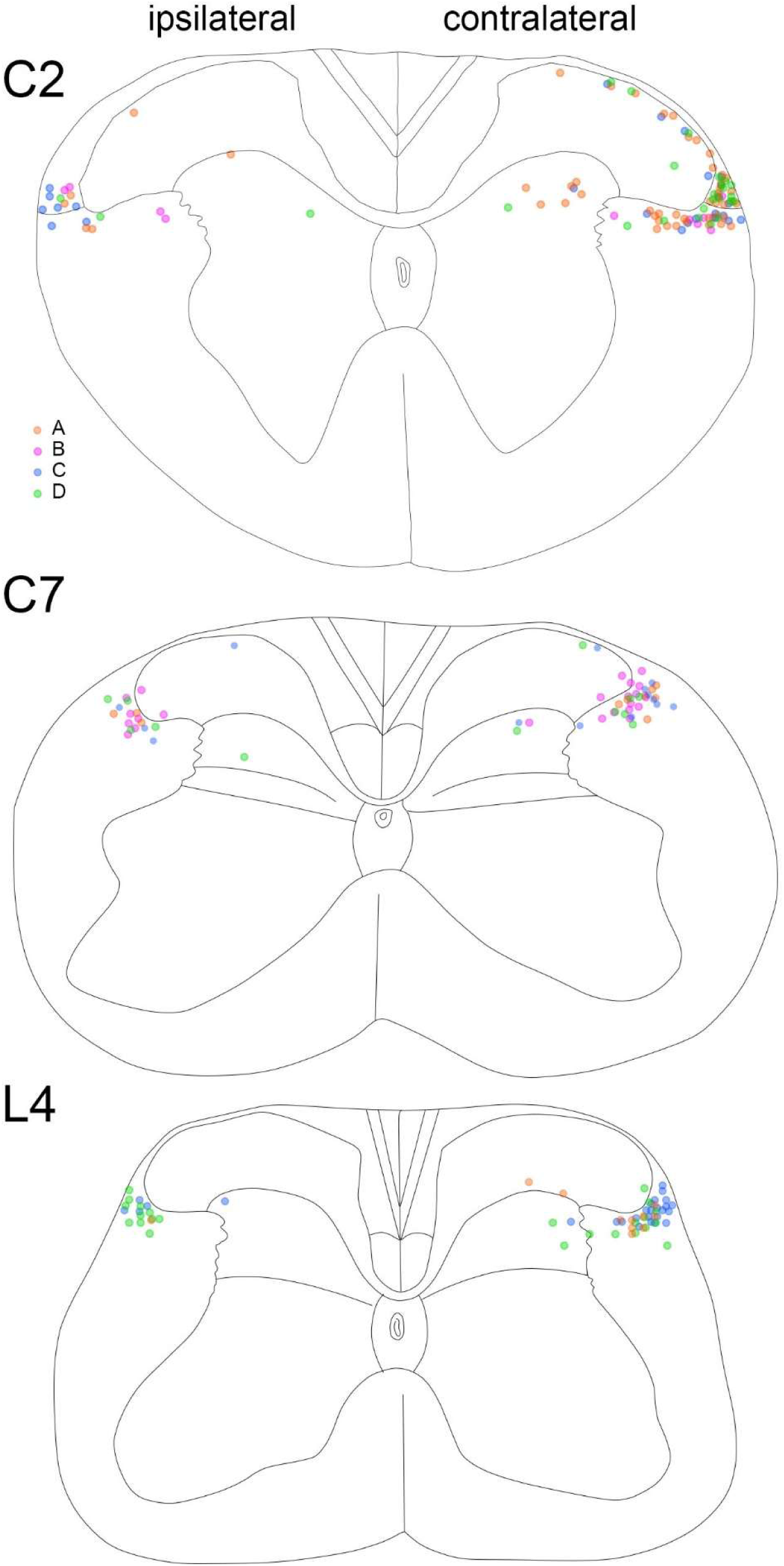
The distribution of retrogradely labelled cells in the C2, C7 and L4 segments from the experiments in which the CTB injection site was restricted to the superior colliculus. Cells from each mouse are represented by a different colour, with orange, pink, blue and green circles showing cells from mice with injections illustrated in parts A, B, C and D of Fig S3. The cells from one animal (pink circles) are not shown on the L4 plot, as they are illustrated in Fig 3. Spinal cord outlines were taken from the Allen Brain Atlas (https://mousespinal.brain-map.org/imageseries/showref.html). Quantitative data from these experiments are provided in Table 1.

In the C2 segment, the LSN is displaced ventrally by the LCN. These two nuclei can readily be distinguished based on the presence of axons containing neuropeptides (including substance P, dynorphin, enkephalin and somatostatin), which are present in the LSN^48–50^, but not the LCN^49^. We therefore used immunohistochemical detection of substance P to differentiate between LSN and LCN (Figure S4). As with the L4 and C7 segments, retrogradely labelled cells were concentrated in the LSN, particularly on the contralateral side. A major difference between the C2 segment and those of the enlargements (L4 and C7) was the presence of many retrogradely labelled cells in the LCN, and these were more numerous on the contralateral side (Table 1, Figure 5). However, we also found considerably higher numbers of cells in lamina I on the contralateral side. If we excluded cells in the LCN, retrogradely labelled cells in LSN therefore accounted for a much lower proportion of the labelled neurons (54%) than was the case for L4 and C7 (Table 1).

Calbindin is known to be present in many LSN projection neurons in the rat^43^, and our finding of input to the SC when intraspinal injections of AAV1.CAG.Cre^ON^.tdTomato extended into the LSN in *Calb1*^Cre^ mice indicates that at least some spinotectal neurons in the LSN express calbindin. To confirm this, we carried out immunohistochemistry for calbindin on transverse sections through the L3 segments of the 4 mice that had received injections of CTB that were restricted to the SC. We identified between 12 and 32 retrogradely labelled neurons in the contralateral or ipsilateral LSN in these sections, and found that 83% (range 69-90%) of these were calbindin-immunoreactive (Figure 6).

### Synaptic input to spinotectal LSN neurons

The LSN receives little or no input from primary afferents^48,50,51^, but is innervated by at least three molecularly-distinct classes of superficial dorsal horn excitatory interneuron, those expressing substance P, neuropeptide FF (NPFF) or the gastrin-releasing peptide receptor (GRPR)^52–55^.

However, the proportions of excitatory synapses accounted for by each of these cell types is not known. We therefore began by estimating the proportions of glutamatergic boutons that could have originated from each of these sources. Glutamatergic boutons in the LSN contain VGLUT2^56–58^, and we examined transverse sections from two *Grpr*^CreERT2^;Ai9 mice that had been immunoreacted to reveal tdTomato, substance P, pro-NPFF (the precursor for NPFF^53^) and VGLUT2 (Figure S5).

Although tdTomato-labelled axons (derived from GRPR-expressing neurons) and boutons containing substance P or pro-NPFF were present throughout the LSN, we observed that the density of tdTomato-labelled axons was considerably higher in the lateral part of the nucleus, compared to the medial part, and we therefore divided the LSN into lateral and medial parts for the purpose of this analysis (Figure S5A). Some GRPR-expressing dorsal horn neurons are known to contain *Tac1* mRNA, which encodes preprotachykinin 1, the precursor for substance P^54^. However, there is little co-expression of *Tac1* and *Npff*^53^, and axons of neurons that express GRPR very seldom contain pro-NPFF^59^. Consistent with this, we found that some tdTomato-labelled axonal boutons in the LSN were substance P-immunoreactive, but boutons containing both substance P and pro-NPFF were extremely rare, and no overlap was seen between tdTomato and pro-NPFF (Table S1, Figure S5B- D). Overall, immunoreactivity for tdTomato, substance P and/or pro-NPFF was found in 46% of VGLUT2 boutons in the LSN (Table S1). tdTomato-labelled boutons (originating from GRPR-expressing cells) accounted for 19% of LSN VGLUT2-positive boutons, while substance P was seen in 22%, and pro-NPFF in 11%.

To determine whether any of these types of bouton were presynaptic to spinotectal neurons in the LSN, we used retrograde transport of AAV11.CAG.tdTomato or AAV11.CAG.GFP, injected into the superior colliculus of 2 wild-type mice and 2 *Grpr*^CreERT2^;Ai9 mice (respectively), to provide extensive labelling of the dendritic trees of these cells. In preliminary experiments, we found that it was difficult to visualise injection sites with this approach, due to the widespread expression of the fluorescent protein by local neurons, and those with axons entering the region. We therefore included a small amount of cholera toxin B subunit conjugated to either Alexa 488 or Alexa 647 in the injectate, to allow accurate visualisation of the centre of the injection site^60^, which was located in the SC in all cases (Figure S6). Within the spinal cord, neurons retrogradely labelled with tdTomato or GFP were present within the LSN, but also in other regions, likely reflecting uptake from structures outside the SC, such as the PAG. Nonetheless, at least some of the retrogradely labelled neurons in LSN presumably projected to the SC.

To look for input from either substance P- or NPFF-expressing boutons onto retrogradely labelled LSN neurons, we immunoreacted sections from the lumbar enlargement of the 2 wild-type mice to reveal tdTomato, VGLUT2, and PSD95, together with either substance P or pro-NPFF. We identified a mean of 162 PSD95 puncta (range 53-679) on 10 tdTomato-labelled LSN neurons (5 from each mouse), and found that 13.7% (2.9-21.5%) of these were associated with boutons immunoreactive for substance P and VGLUT2 (Figures 7, S7). We observed that synapses from pro-NPFF-labelled boutons onto retrogradely labelled LSN neurons were extremely rare, and we therefore did not quantify these. We estimated the proportion of excitatory synaptic input originating from GRPR-expressing cells by immunostaining equivalent sections from the two *Grpr*^CreERT2^;Ai9 mice to reveal GFP, tdTomato and PSD95. We identified a mean of 75 PSD95 puncta (range 46-134) on 10 GFP-labelled LSN neurons (5 from each mouse) and found that 11.5% of these (0-28.4%) were associated with a tdTomato-positive bouton (Figures 7, S7).

**Fig 6.**
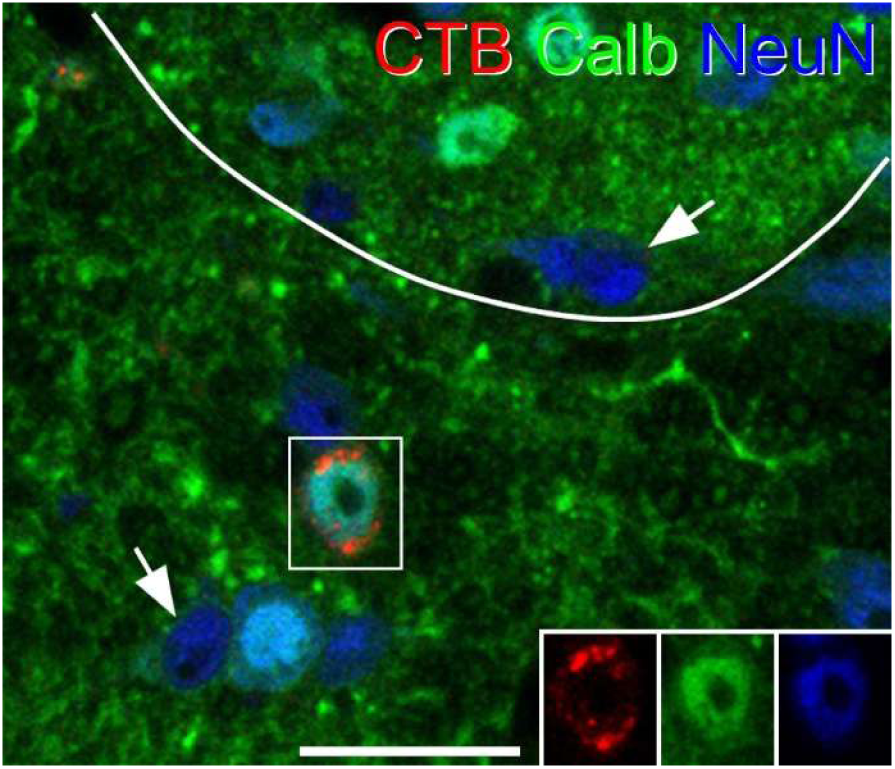
Calbindin expression in a LSN neuron retrogradely labelled from the superior colliculus. The confocal image shows a single optical plane taken from a transverse section through the L3 segment from one of the mice that received an injection of CTB that was restricted to the superior colliculus (illustrated in Fig S3B). The section has been reacted to reveal CTB (red), calbindin (green) and NeuN (blue). A single CTB-positive cell in the LSN is present in the area marked by the box, and this is immunoreactive for calbindin. Two cells that are NeuN-positive but lack calbindin (i.e. calbindin-negative neurons) are indicated with arrows. The line indicates the border between the grey matter and the LSN. The inset shows each type of immunoreactivity separately in the region corresponding to the box. Scale bar = 20 µm.

**Fig 7.**
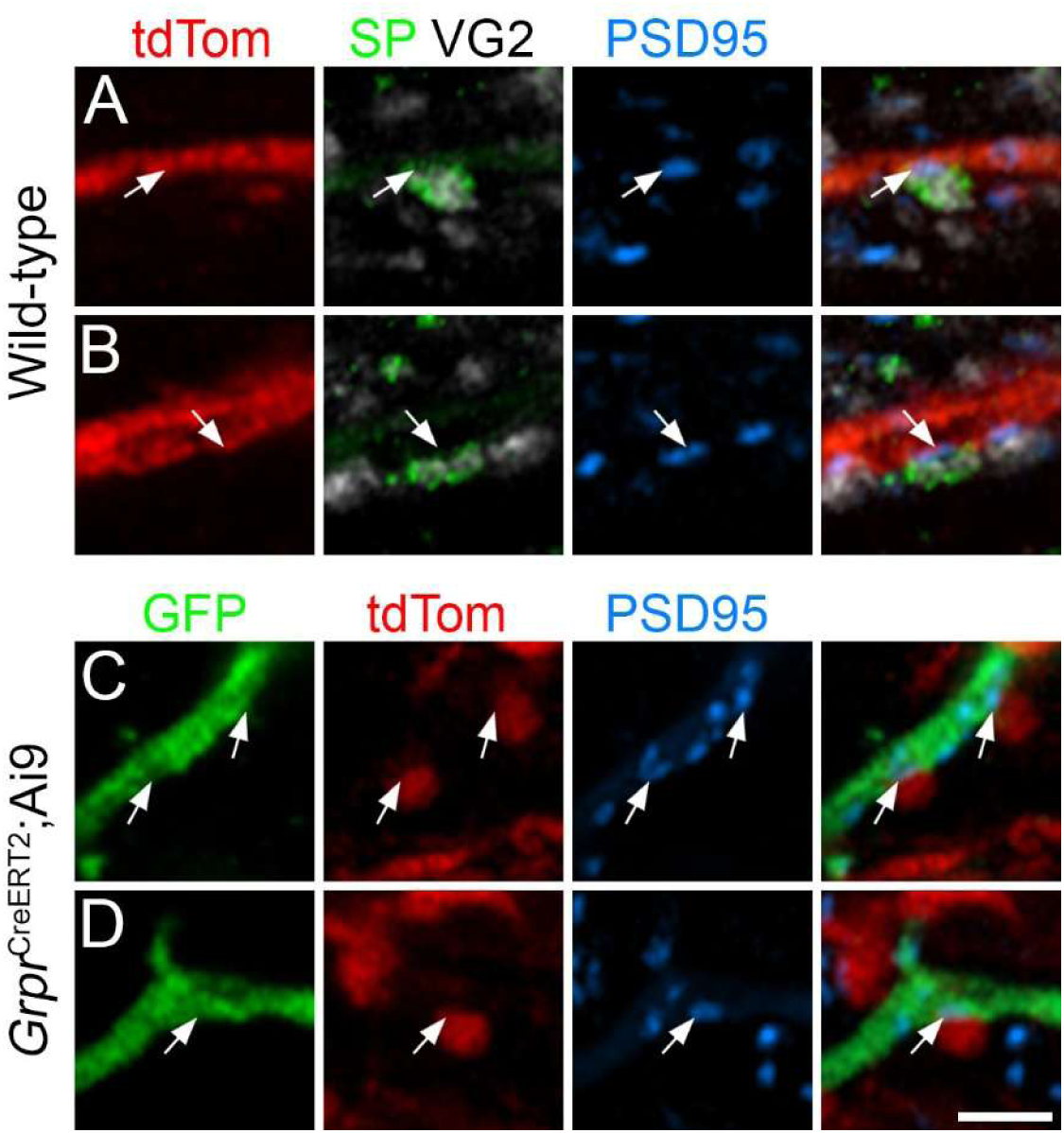
Synaptic input to LSN neurons retrogradely labelled from the superior colliculus in a wild-type (**A**, **B**) or a *Grpr*^CreERT2^;Ai9 (**C**, **D**) mouse. Each row shows a single confocal optical plane through part of the dendritic tree of a retrogradely labelled LSN neuron. **A** and **B** were scanned to reveal tdTomato (tdTom, red), VGLUT2 (VG2, grey), substance P (SP, green) and PSD95 (blue), and include parts of a dendrite belonging to a neuron retrogradely labelled following injection of AAV11.CAG.tdTomato into the superior colliculus. In each row a bouton that is immunoreactive for both VGLUT2 and substance P is in contact with a PSD95 punctum on a dendrite belonging to the cell (arrows). **C** and **D** were scanned to reveal GFP (green), tdTomato (tdTom, red) and PSD95 (blue) and show parts of two dendrites of a neuron retrogradely labelled following injection of AAV11.CAG.GFP into the superior colliculus. Arrows indicate PSD95 puncta on the labelled dendrites that are in contact with tdTomato-positive boutons, which originate from GRPR-expressing neruons. Scale bar = 2 µm. The locations of these synapses on the retrogradely labelled neurons are shown in Figure S7.

## DISCUSSION

The main findings of this study are: (1) that input from the spinal enlargements is largely restricted to the caudal half of the superior colliculus and densest in its lateral part, is associated with the clusters of Pitx2-positive cells located in the intermediate grey layer, and shows a somatotopic organisation, (2) that within the spinal enlargements this input originates mainly from neurons in the LSN, most of which express calbindin, and (3) that LSN neurons projecting to the SC are likely to be innervated by substance P- and GRPR-expressing excitatory interneurons in the superficial dorsal horn.

### Spinal input to the superior colliculus

Our finding of a patchy input from the spinal cord to the intermediate layer of the caudal part of the SC in the anterograde tracing experiments matches those of several previous studies carried out in various species^14–16,38,47^. However, little was known about the cells that give rise to this input. In early retrograde tracing studies, injection sites either included other areas that received spinal input (in particular the PAG)^37–39,42^ or else used the relatively insensitive tracer horseradish peroxidase (HRP)^40,41^. In two studies that restricted HRP injection sites to the SC, numerous retrogradely labelled cells were seen in the LCN, in agreement with our findings, but cells in the lumbar spinal cord were apparently very sparse, and were limited to the deep dorsal horn^40,41^. By restricting injections of the highly sensitive tracer CTB to the SC, we demonstrate that the great majority of neurons in cervical and lumbar enlargements that project to the SC are located in the LSN. Together with the finding that most of these cells contain calbindin, this is consistent with our previous observation that anterograde labelling in SC is seen following intraspinal injection of AAV encoding Cre-dependent tdTomato in *Calb1*^Cre^ mice only when injection sites included the LSN^28^.

Ding et al^13^ recently used fMOST to generate detailed reconstructions of spinal projection neurons, allowing them to assign these to 19 clusters based on their projection patterns. Several of these clusters included cells that projected to the SC, with some preferentially innervating posterior (caudal), lateral or medial parts of the colliculus. Our anterograde tracing studies show that the densest input from the spinal enlargements targeted caudal and lateral parts of the SC, and in the study of Ding et al the clusters with the highest numbers of neurons projecting to these regions were S8 and S11, both of which included many cells in the LSN. This therefore fits with our observations from the retrograde tracing studies. However, neuronal reconstructions performed by Ding et al were restricted to upper cervical segments (C1-C4), and our finding of differences between the distribution of retrogradely labelled cells in C2 compared to C7 and L4 segments suggests that projection patterns may differ between upper cervical segments and the remainder of the spinal cord. Interestingly, most of the clusters that contained neurons projecting to the medial part of the SC in the study of Ding et al included cells located in the dorsal white matter, which may represent a caudal extension of the gracile nucleus^13^. This is consistent with the report that inputs from the gracile nucleus target the medial part of the SC in the rat^11^.

### Neurons in the lateral spinal nucleus

Previous studies have demonstrated that LSN neurons can be retrogradely labelled from several sites in the brain, including the NTS, medullary reticular formation, LPB, PAG, thalamus and hypothalamus^37,42,43,61–71^. This raises the question of whether those cells projecting to the SC represent a distinct subset among LSN projection neurons, or whether the input to SC arises from collaterals of axons that project to other brain regions. Both the S8 and S11 populations of Ding et al^13^ have several other targets in brainstem and thalamus, suggesting that the projections to SC are collaterals from cells that also project to other sites.

Relatively few studies have investigated the physiological properties of LSN neurons. Menétrey et al^62^ reported that most lumbar LSN projection neurons were not spontaneously active, and had poorly defined receptive fields on the caudal part of the body that responded inconsistently to mechanical stimuli applied to skin or deep tissues. Sikandar et al^51^ found that neurons in LSN could be activated by strong mechanical stimuli that were thought to excite receptors in deep tissues, but not by heat stimuli applied to skin. They also reported sensitisation of these cells following acute muscle inflammation, with development of sensitivity to brush. However, other studies have demonstrated Fos expression in the LSN after noxious heating of the skin^72,73^, and a recent paper reported that some LSN neurons responded to intradermal injection of chloroquine into the nape of the neck, suggesting a role in non-histaminergic itch^74^.

The LSN receives few, if any, direct projections from primary afferents^51^, although electrophysiological studies have revealed some monosynaptic input from both Aδ and C fibres^75,76^, which may be through synapses located on dendrites that extend outside the nucleus. We demonstrate here that nearly half of the excitatory synapses within the LSN originate from neurons that express substance P, NPFF or GRPR, which are likely to be located in the superficial laminae^52,53,55^. We also show that LSN neurons retrogradely labelled from injections of AAV targeted on the SC receive excitatory synapses from both the substance P and GRPR-expressing subsets, many of which are known to respond to noxious mechanical and thermal, as well as pruritic, stimuli^54,77^. Consistent with this, we previously reported that chemogenetic activation of GRPR-expressing dorsal horn neurons resulted in Fos expression by many projection neurons in the LSN^55^. Since GRPR-expressing neurons in the superficial dorsal horn are strongly implicated in itch^78–80^, their input is likely to underlie responses of LSN neurons to pruritic stimuli^74^.

### A role for LSN cells in head movement directed towards a noxious or pruritic stimulus

Since neurons in the LSN can be activated by noxious stimuli, those projecting to SC are likely to be an important source of nociceptive input to this structure. Cells in the SC that respond to noxious stimuli are located in the deeper layers^5,6,8,9^, in accordance with our finding that spinal inputs target the SGI. SC cells responding to noxious stimuli often have large receptive fields that can extend over the whole body, but with strongest input from a particular body region^5,6,8^, and this is consistent with the partially overlapping pattern of input that we see following anterograde tracing from both lumbar and cervical enlargements. Cells responding to low-threshold input in the SC show a clear somatotopic pattern with face, forelimb and hindlimb represented in rostral, central and caudal parts of the SC^9^. However, the evidence for a somatotopic pattern for nociceptive neurons is less clear, since in one study these were concentrated in the rostral SC^9^, while two other studies identified many in the central and caudal parts of the colliculus^6,8^. It is possible that complications due to the large receptive fields and the effects of general anaesthesia on responses to noxious stimuli in *in vivo* recordings account for the lack of evidence for a clear somatotopic pattern in physiological studies. LSN neurons are unlikely to account for all of the nociceptive input to SC, since it is also innervated by cells in the LCN, many of which respond to noxious stimuli^81,82^.

However, although the LCN is somatotopically organised^81,83^, it is not known whether this organisation is maintained in its projection to the SC.

The SC drives coordinated head movements in response to a wide variety of sensory stimuli^2–4^.

Two main behavioural patterns are recognised: orientation towards a stimulus and defence-like behaviours, and these are mediated by crossed and uncrossed descending pathways, respectively^84^. Direct evidence that the SC is responsible for movements of the head towards a noxious stimulus was provided by Wang and Redgrave^85^, who showed that injection of the GABA_A_ receptor agonist muscimol into the SC reduced biting and licking of the affected hindlimb in the formalin test. In addition, lesions to the contralateral output from the SC were found to increase the latency for removal of a clip applied to the hindpaw^86^.

Our finding of an association between spinotectal axons and clusters of Pitx2-expressing cells suggests that these axons target the Pitx2 cells, either directly through synapses on their dorsal dendrites^45^, or indirectly via neurons that are presynaptic to these cells (Pitx2^ON-Pre^ cells), which are found throughout the SGI^87^. Since activation of Pitx2 cells can induce turning of the head^45^, this pathway is likely to contribute to the attention to a painful or injured limb to allow licking.

Interestingly, the densest input from the spinal enlargements was to the extreme lateral part of the SC. This has been identified as a specific zone, known as SC.l, and is thought to be responsible for head-turning^2^, as well as for oro-facial movements such as licking and biting during food consumption^88^. It may therefore also underlie site-directed licking in response to a noxious stimulus applied to the limbs^36^.

To the best of our knowledge, little or nothing is known about a role for the SC in itch. However, our finding of input from GRPR-expressing interneurons to LSN neurons, including those that project to the SC, together with the report that some LSN neurons upregulate Fos after a pruritic stimulus^74^, raises the possibility that the SC also receives pruriceptive input. It may therefore co- ordinate movement of the head to allow actions that relieve itch, such as biting of the affected skin on the hindlimb^36,89^.

We recently identified 5 transcriptomic classes among those ALS neurons that are derived from the Phox2a lineage^29^, allowing us to define functional populations among these cells^23,28^. However, LSN neurons will have been substantially under-represented in this dataset, due to the relatively infrequent capture of these cells in the Phox2a::Cre mouse that we used^90^. A more complete analysis of ALS neurons is likely to reveal molecular markers for LSN populations and therefore allow targeting of those that project to the SC, in order to determine their role in movements directed towards an area of injury.

## METHODS

### Animals

All experiments were approved by the Ethical Review Process Applications Panel of the University of Glasgow, and were performed in accordance with the European Community directive 86/609/EC and the UK Animals (Scientific Procedures) Act 1986. The study was carried out in compliance with the ARRIVE guidelines. Mice of both sexes were used in this study. They were aged 4 to 11 weeks and weighed between 16 and 31 g at the time of perfusion or at the start of any surgical procedures.

### Surgery

All surgical procedures involving intraspinal or intracranial injections were carried out while the mice were anaesthetised with isoflurane (1-2%), and all of these animals received perioperative analgesia (buprenorphine 0.1 mg/kg and carprofen 10 mg/kg, s.c.).

Perfusion fixation was performed following induction of deep anaesthesia with pentobarbitone (20 mg, i.p.). A brief rinse with Ringer’s solution was followed by perfusion with 250 ml of fixative, which contained 4% freshly depolymerised formaldehyde in phosphate buffer. For experiments involving *in situ* hybridisation, the fixative was made up in RNAse-free buffer. Spinal cords and brains were dissected out and stored in the same fixative for 2 hours.

Details of the intraspinal injections of AAV1.CAG.Cre^ON^.tdTomato into *Calb1*^Cre^ mice that are illustrated in Figures 1 and S1 are provided in a previous paper^28^.

For combined anterograde tracing and fluorescence *in situ* hybridisation, 3 female *Calb1*^Cre^ mice received intraspinal injections of AAV1.CAG.Cre^ON^.tdTomato (University of Zurich, Viral Vector Facility v167-1) into the L3, L4 and L5 segments on the right side, and these were performed as described previously^28^. Briefly, animals were placed in a stereotaxic frame, and the T12 and L1 vertebrae were exposed and clamped. Injections were made into the L3 and L5 segments through the intervertebral spaces on either side of the T13 vertebra, while those into the L4 segment were made through a small hole drilled through the lamina of T13. The injections were made 400 µm lateral to the midline and 300 µm deep to the spinal cord surface, using glass micropipettes (outer diameter ∼60 µm) attached to a 10 µL Hamilton syringe. At each site, a volume of 300 nL containing 9.48 × 10^7^ gene copies was injected at a flow rate of 30 nL per minute, and after each injection the pipette was left in place for 5 minutes to minimise leakage back up the track. The mice survived for between 46-54 days and were then deeply anaesthetised and perfused with fixative.

Four wild-type mice (two of each sex) received intraspinal injections of AAV9.CB7.mCherry (Addgene 105544; 6.9 × 10^8^ gene copies in 300 nl) into the C8 segment and AAV9.CB7.GFP (Addgene 105542; 4.5 × 10^8^ gene copies in 300 nl) into the L3 segment, both on the right side. The L3 injection was carried out as described above. For the C8 injections, the head was held in place with ear bars in a stereotaxic frame and the T2 spine was clamped. Injections were made (as described above) in the space between the C6 and C7 vertebrae. The mice survived for between 47 and 57 days and were then deeply anaesthetised and perfused with fixative.

Injections of CTB that were restricted to the SC were made in 4 wild-type mice (one male, three female). The animals were placed in a stereotaxic frame, the skull was exposed, and injections consisting were made approximately 4.4 mm caudal to Bregma, 0.8 mm to the left of the midline and 0.8 mm below the brain surface through a glass micropipette (outer diameter ∼60 µm) attached to a PV 800 Pneumatic Picopump (WPI). The pipette was left in place for at least 5 minutes. The mice survived for 3-4 days before perfusion fixation under terminal anaesthesia.

To investigate synaptic inputs to spinotectal LSN cells, we injected: (1) 200 nl of a mixture containing AAV11.CAG.tdTomato (BrainCase BC-0868, 8.0 × 10^8^ GC) and CTB conjugated to Alexa 488 (0.2 or 0.05%) into the SC of two male wild-type mice; and (2) 200 nl of a mixture containing AAV11.CAG.GFP (BrainCase BC-0251, 8.16 × 10^8^ GC) and CTB conjugated to Alexa 647 (0.05%) into the SC of two male *Grpr*^CreERT2^;Ai9 mice, which had been treated with tamoxifen, as described previously^80^. Injections were performed as described above. After a two week survival time, the mice were perfused with fixative. Brain sections were cut transversely to visualise the centre of the injection site, which was revealed with either Alexa 488 or Alexa 647. Sagittal sections of spinal cord were cut from the side contralateral to the brain injection from the L2-L5 segments and processed for immunohistochemistry.

### Tissue processing and image analysis

Coronal sections of brain (16 or 50 µm thick) and transverse or parasagittal sections of spinal cord (60 or 30 µm thick) were cut with a vibrating blade microtome (Leica VT1200) or a cryostat (Leica CM1950). Multiple-labelling immunofluorescence reactions were carried out on free-floating sections as described previously^23,54^. Briefly, sections were incubated for 1-3 days at 4 °C in primary antibodies diluted in phosphate buffered saline (PBS) that contained 0.3M NaCl, 0.3% Triton X-100 and 5% normal donkey serum, and then for 2-18 h in appropriate species-specific secondary antibodies that were raised in donkey and conjugated to Alexa 488 or Alexa 647 (from Jackson Immunoresearch) or Alexa 555-plus (Thermo Fisher Scientific). In some cases, sections underwent dehydration and rehydration in graded methanol/water mixtures (5-10 mins each in 20%, 40%, 60%, 80%, 100%, 80%, 60%, 40%, 20% methanol) in order to enhance antibody penetration. A list of primary antibodies used in the study is provided in Table S2. Sections were scanned with either a Zeiss LSM710 confocal microscope with Argon multi-line, 405 nm diode, 561 nm solid state and 633 nm HeNe lasers, or with a Zeiss LSM900 Airyscan confocal microscope with 405, 488, 561 and 640 nm diode lasers. Confocal image stacks were obtained through a 20x dry lens (numerical aperture, NA, 0.8), or through 40x (NA 1.3) or 63x (NA 1.4) oil immersion lenses with the confocal aperture set to 1 Airy unit or less.

Combined fluorescent *in situ* hybridisation and immunohistochemistry was performed on sections through the superior colliculus from 3 female *Calb1*^Cre^ mice that had received injections of AAV1.CAG.Cre^ON^.tdTomato into the L3, L4 and L5 spinal segments on the right side 46-54 days prior to perfusion fixation. Coronal sections (16 µm thick) were slide-mounted and *in situ* hybridisation was carried out using a RNAscope multiplex fluorescent kit (v2) according to the manufacturer’s instructions, with a probe against *Pitx2*, which was revealed with TSA Vivid 650. Slides were incubated overnight in antibody against mCherry, which was revealed with secondary antibody conjugated to Alexa 555-plus.

To directly compare lumbar and cervical inputs to the SC, we immunoreacted sections from mice in which AAVs coding for GFP and mCherry had been injected into the two spinal enlargements.

Coronal sections (50 µm thick) were immunoreacted to reveal GFP and mCherry, and scans were obtained from different rostrocaudal levels through the colliculus.

Immunoperoxidase labelling of CTB brain injection sites was carried out by incubating sections in anti-CTB at 1:100,000 overnight, followed by incubation in biotinylated secondary antibody and avidin-peroxidase, which was revealed with diaminobenzidine in the presence of H_2_O_2_. For the analysis of retrograde labelling in these experiments, we analysed between 7 and 19 transverse sections (60 µm thick) from C2, C7 and L4 segments that had been immunoreacted for CTB and NeuN. Cells were included in the analysis if they were entirely contained within the section, or if part of the soma was present at the top surface, but were excluded if part of the soma was present at the bottom of the section (i.e. if part of the soma appeared on the last scan in the z-series).

Retrogradely labelled cells were plotted with Neurolucida, and these plots were then morphed onto a standard outline of the corresponding segment in Inkscape.

To determine the proportion of LSN cells retrogradely labelled from the SC that contained calbindin, we immunostained 30 µm thick transverse sections from the L3 segments of the 4 mice that had CTB injections restricted to the SC. Sections were reacted with antibodies against CTB and calbindin, as well as the monoclonal NeuN antibody, which labels all neurons in the spinal cord^91^. Scans were analysed with Neurolucida and the presence or absence of calbindin-immunoreactivity in each CTB-labelled LSN neuron was noted. In cases of weak immunoreactivity, the luminance value for calbindin immunoreactivity over the cell was compared to that of the 5 closest neurons identified as non-immunoreactive, and if this was more than 4 times the mean value over those cells, the LSN neuron was defined as calbindin-immunoreactive. To avoid double-counting of neurons, sections were all mounted in the same orientation, and LSN neurons for which part of the nucleus was cut at the lower face of the section were excluded from the analysis.

We estimated the proportions of glutamatergic (VGLUT2-positive) boutons in the LSN that contained tdTomato, substance P or pro-NPFF in two male *Grpr*^CreERT2^;Ai9 mice, which had been treated with tamoxifen as described previously^80^. A single 60 µm thick transverse section through the L4 segment of each animal was scanned and analysed using Neurolucida software. The area corresponding the LSN was initially defined and divided into medial and lateral parts, as described in the Results section. A 3 × 3 µm grid was placed over the area and a single z-section was selected from each animal. Working through the grid squares (from top left to bottom right) with only the VGLUT2 channel visible, we identified the VGLUT2-immunoreactive bouton that was closest to the bottom right corner of each square. The other channels were then viewed and the presence or absence of immunostaining for tdTomato, substance P and pro-NPFF was noted.

Excitatory synaptic input from substance P- or GRPR-expressing boutons were analysed using Neurolucida software. Cell bodies and parts of the dendritic trees of the neurons were initially reconstructed from 60 µm thick sagittal sections through the LSN, and the locations of PSD95 puncta on these cells were plotted, with only fluorescent protein and PSD95 channels visible. The remaining channels were then viewed and the presence or absence of immunostaining adjacent to the PSD95 puncta was noted.

## ACKNOWLEDGEMENTS

We thank Iain Plenderleith for expert technical assistance, Dr YanGang Sun for the gift of GRPR^CreERT2^ mice Dr Ana Gonzalez-Rueda for helpful discussion. Financial support from the Wellcome Trust (Grant numbers 219433/Z/19/Z and 304005/Z/23/Z) and the Biotechnology and Biological Sciences Research Council (Grant number UKRI2956) is gratefully acknowledged. For the purpose of open access, the author has applied a Creative Commons Attribution (CC BY) licence to any Author Accepted Manuscript version arising from this submission.

## AUTHOR CONTRIBUTION

M.A.H., W.M., A.M.B. and A.J.T. conceived the project and designed experiments; M.A.H., M.Y., W.M., A.C.D. and E.P. performed experiments; M.A.H., M.Y. and W.M. analysed data; M.A.H., W.M. and A.J.T. wrote the paper; all of the authors provided feedback and contributed to the editing of the manuscript.

## DATA AVAILABILITY

The datasets generated and analysed during the current study are available from the corresponding author on reasonable request.

