## Supplementary information for "Neurons in the lateral spinal nucleus give rise to the spinotectal component of the anterolateral system in the mouse"

**Supplementary Figures 1-7**

**Supplementary Table 1-2**

**Figure S1**

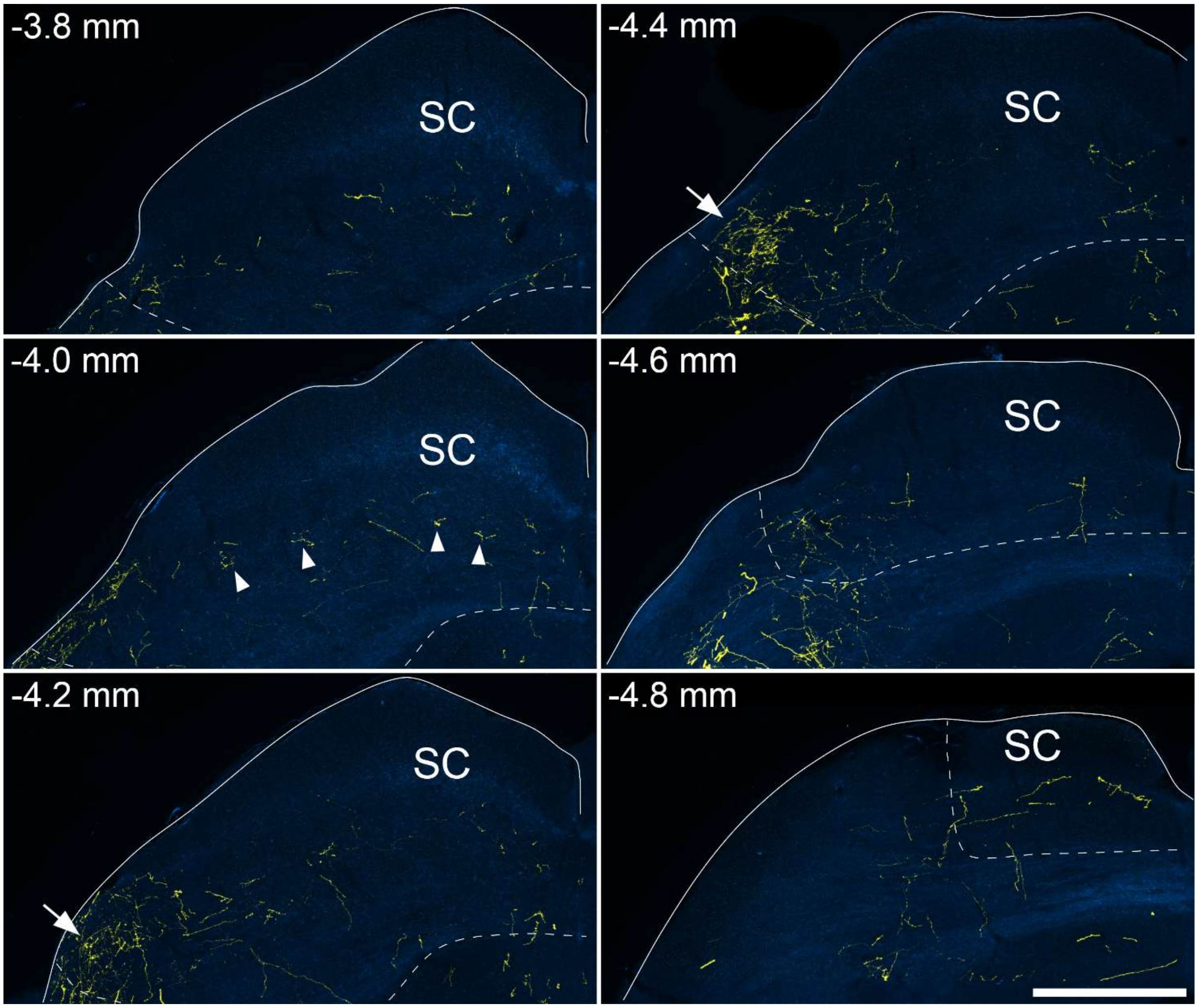


Axonal projections from calbindin-positive neurons in the cervical enlargement to the superior colliculus. Images show labelling with tdTomato (yellow) superimposed on dark field images (blue) at different levels of the contralateral superior colliculus (SC) from a Calb1$^{Cre}$ mouse that had received an intraspinal injection of AAV1.Cre$^{ON}$.tdTomato into the C8 segment on one side. Estimated levels are indicated as millimetres caudal to the bregma and approximate borders of the superior colliculus are shown with dashed lines. Anterograde labelling is present in each of these sections, but is densest in the extreme lateral part of the SC at 4.2 and 4.4 mm caudal to bregma (arrows). At 4 mm caudal to bregma there is some evidence of patchy labelling (arrowheads). A similar pattern of labelling was seen in 2 other *Calb1*$^{Cre}$ mice that had received intraspinal injections of AAV1.Cre$^{ON}$.tdTomato into the C8 segment. Scale bar = 500 μm.

**Figure S2**

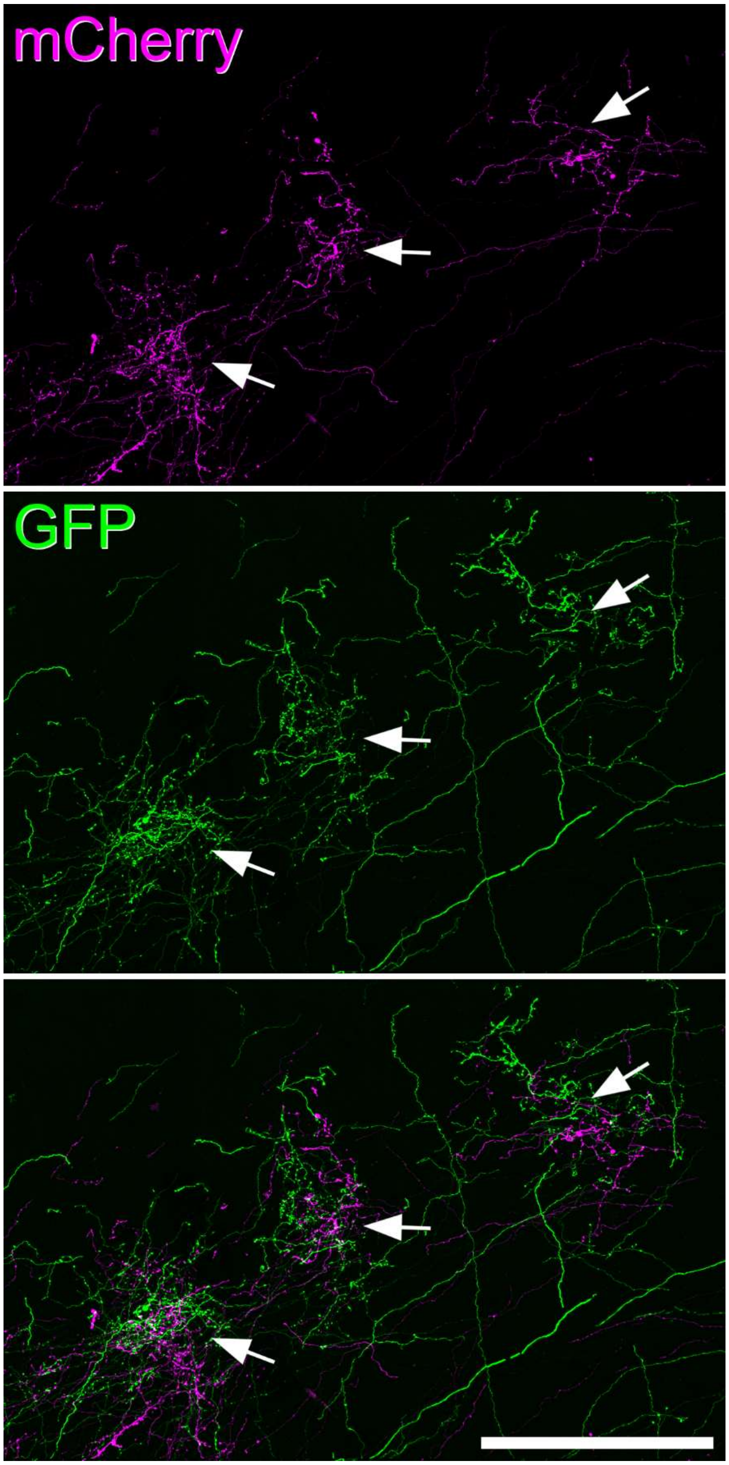


Overlap of axonal projections from lumbar and cervical enlargement to the superior colliculus. Images show axonal labelling with GFP (green) and tdTomato (magenta) in the section illustrated in Figure 3 at the level 4.4 mm caudal to bregma. At this magnification it can be seen that axons from both enlargements are intermingled within clusters in the superior colliculus (arrows). Scale bar = 200 µm.

**Figure S3**

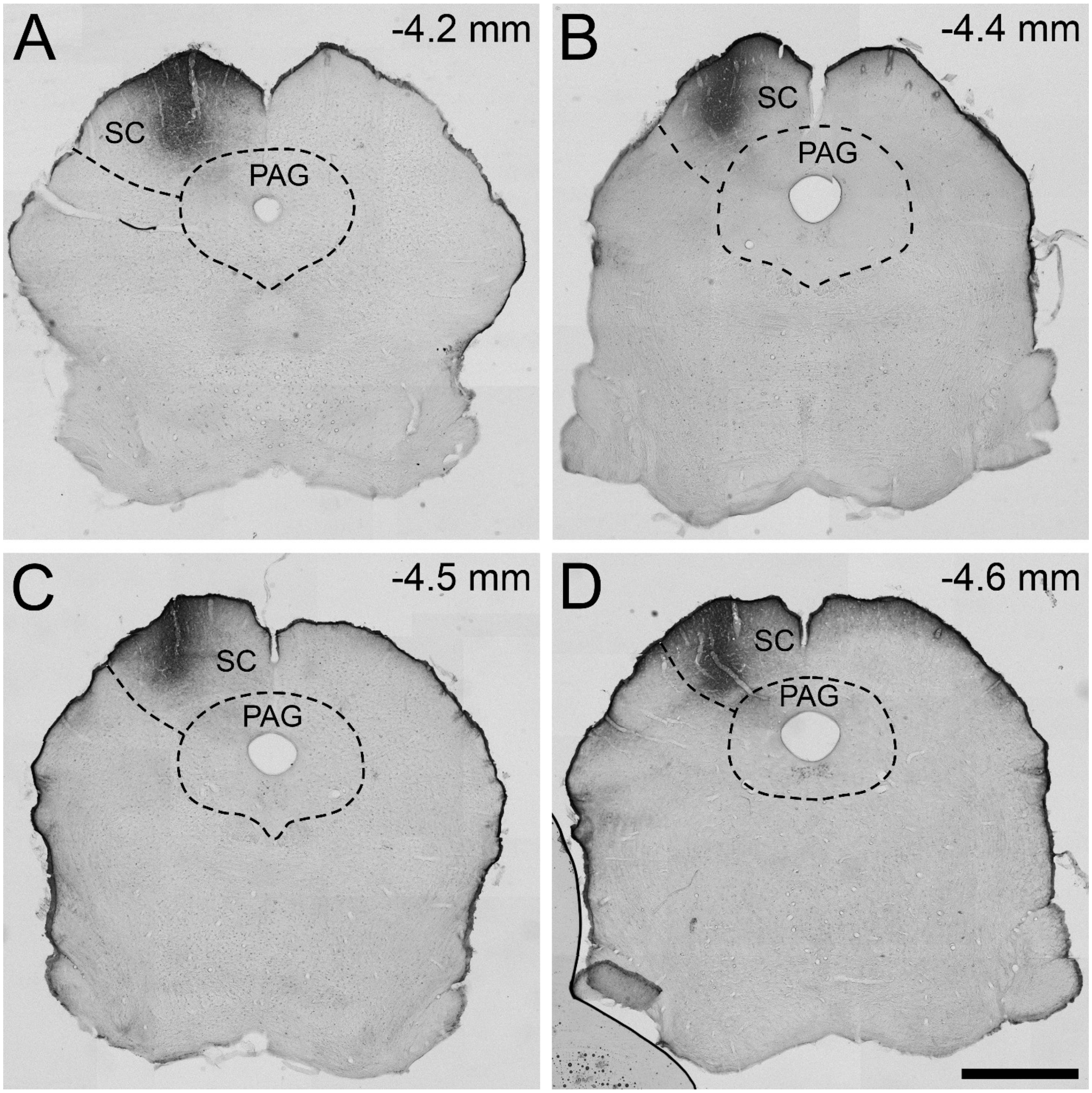


Cholera toxin B subunit (CTB) injection sites in 4 mice in which the injection was restricted to the superior colliculus. Estimated distances caudal to bregma are indicated for each case. SC, superior colliculus; PAG, periaqueductal grey matter. Scale bar = 1 mm.

**Figure S4**

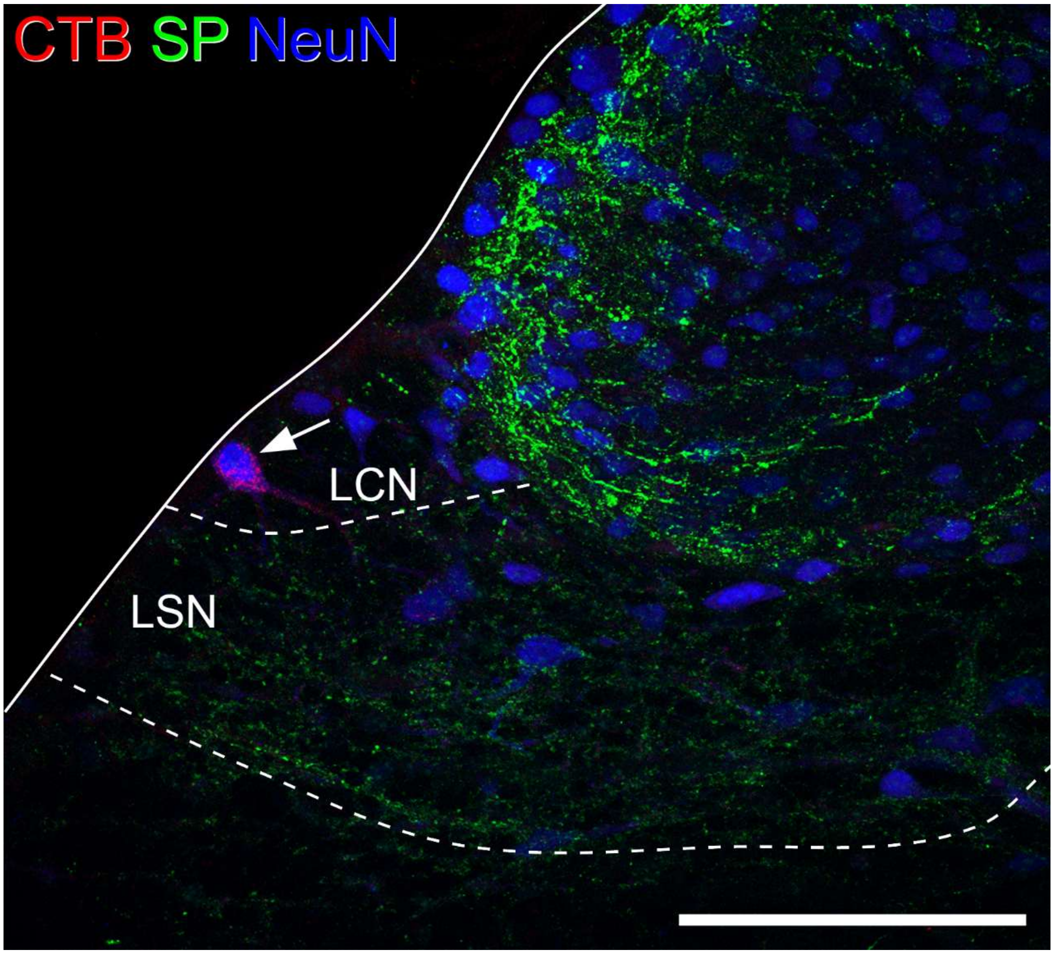


A neuron in the ipsilateral lateral cervical nucleus (LCN) in the C2 segment that was retrogradely labelled from the superior colliculus following the injection of CTB that is illustrated in Fig S3C. The section was immunostained to reveal CTB (red), substance P (SP, green) and NeuN (blue). There is very dense SP labelling in the superficial laminae of the dorsal horn, and SP-immunoreactive axons are also present in the lateral spinal nucleus (LSN). However, lack of SP labelling in the LCN allows this to be distinguished from the LSN. The arrow indicates the retrogradely labelled neuron. Scale bar = 100 µm.

**Figure S5**

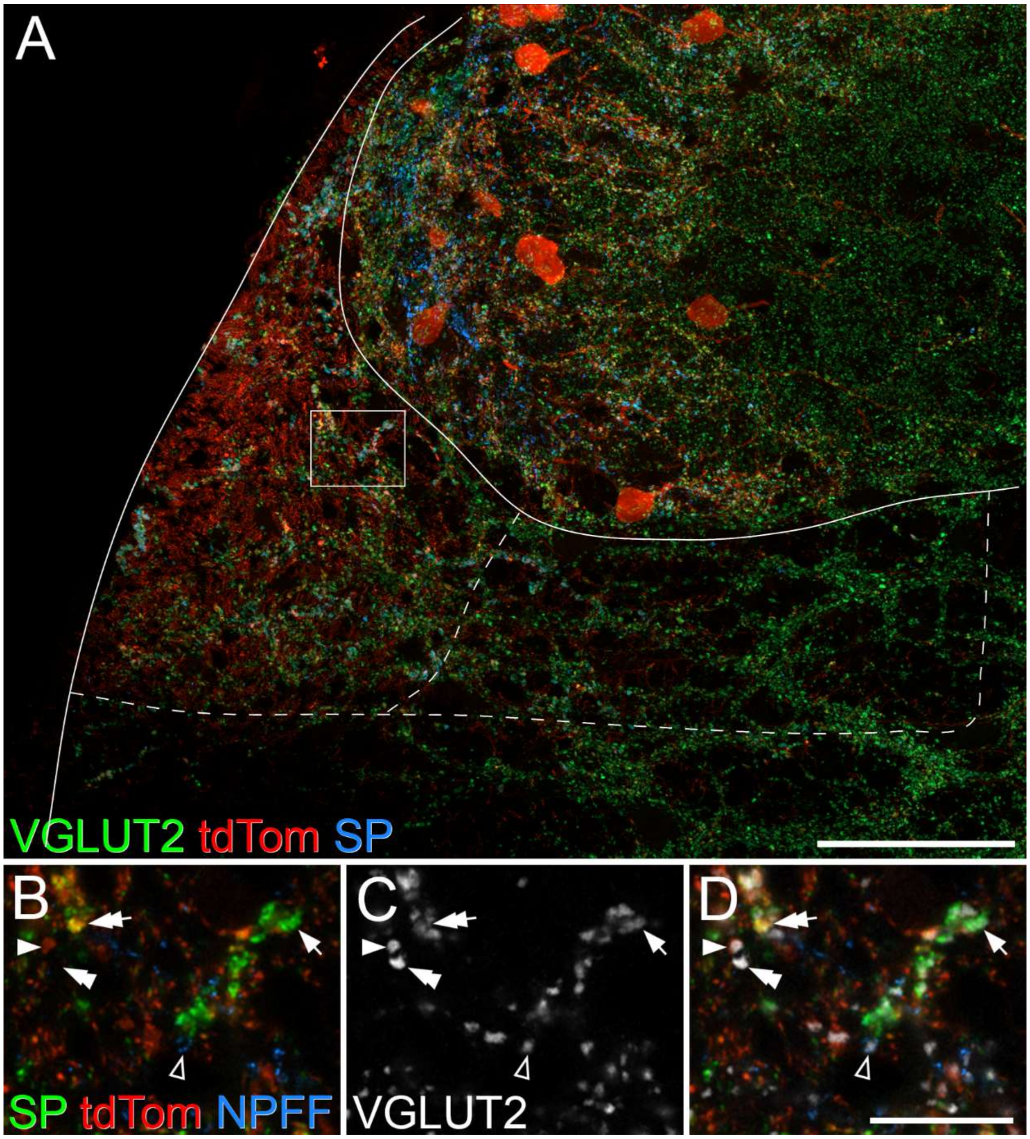


Immunostaining for VGLUT2, tdTomato, substance P and pro-NPFF in a transverse section through the lateral spinal nucleus from a GRPR$^{CreERT2}$;Ai9 mouse. **A**: confocal image showing the extreme lateral part of the dorsal horn and lateral spinal nucleus (LSN) stained to reveal VGLUT2 (green), tdTomato (tdTom, red) and substance P (SP, blue). For the quantitative analysis the LSN was divided into 2 regions (outlined by the dashed line), with the lateral part having a much higher density of tdTomato-labelled profiles. **B**-**D**: high magnification views corresponding to the box in **A**, showing immunostaining for substance P (green), tdTomato (red), pro-NPFF (NPFF, blue) and VGLUT2 (grey). Various types of VGLUT2-positive bouton are identified as follows: single arrow – substance P-positive, double arrow – substance P- and tdTomato-positive, open arrowhead – pro-NPFF-positive, single arrowhead – tdTomato-positive, double arrowhead – negative for each of these markers. Scale bars = 50 µm (A), 10 µm (B-D).

**Figure S6**

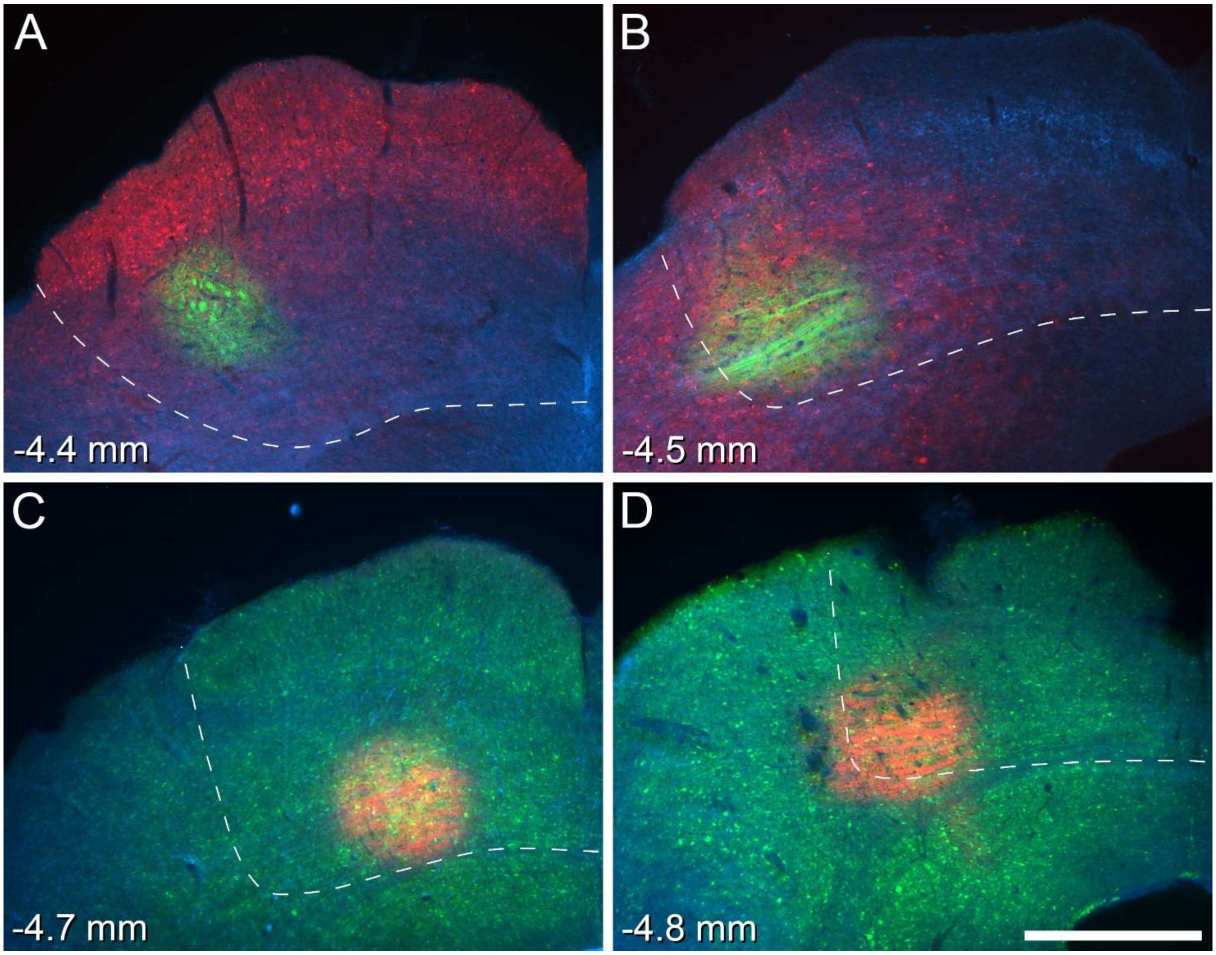


Combined viral and CTB injection sites in 4 mice used to examine synaptic input to retrogradely labelled LSN neurons. **A**,**B**: Injections of AAV11.CAG.tdTomato and CTB-Alexa488 targeted on the superior colliculus in two wild-type mice. The Alexa488 fluorescence (green) shows the CTB labelling, which was used to reveal the centre of the injection site. Expression of tdTomato (red) is far more widespread, and presumably results from uptake by neurons at the centre of the injection, as well as by those with axons extending into this region. Dark-field illumination is shown in blue. **C**,**D**: Injections of AAV11.CAG.GFP and CTB-Alexa647 targeted on the superior colliculus of two $GRPR^{CreERT2}$;Ai9 mice. Alexa647 fluorescence (red) shows CTB labelling at the centre of the injection site, while GFP expression (green) is widespread. Again, dark-field illumination is shown in blue. Estimated levels are indicated as millimetres caudal to the bregma and the approximate boundaries of the superior colliculus are shown with dashed lines. Scale bar = 500 µm.

**Figure S7**

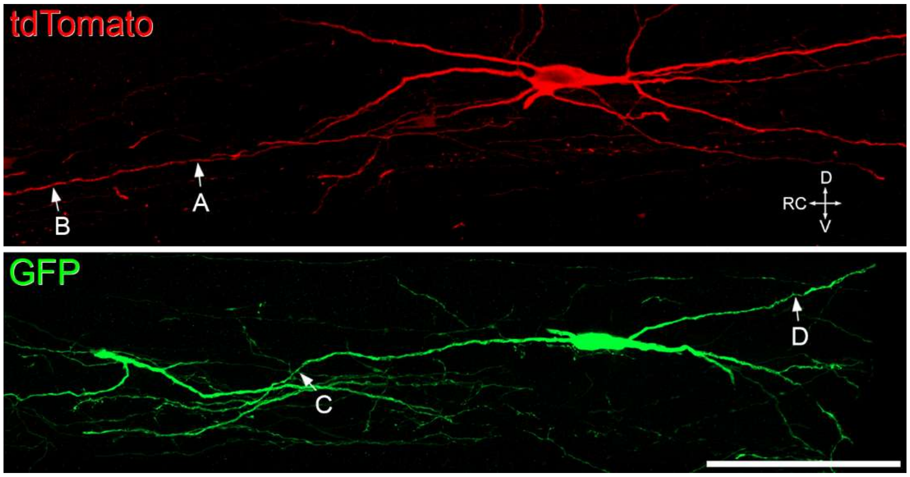


Neurons in the lateral spinal nucleus that were labelled from the superior colliculus and used to investigate synaptic input. The upper panel shows a labelled neuron from a wild-type mouse that had been injected with AAV11.CAG.tdTomato, while the lower panel shows a labelled neuron from a *Grpr*$^{CreERT2}$;Ai9 mouse that had been injected with AAV11.CAG.GFP. A-D show the locations of synapses illustrated in the corresponding parts of Figure 7. Orientation marker: D, dorsal, V, ventral, RC, rostrocaudal. Scale bar = 100 µm. These cells were obtained from the experiments shown in Figure S6B and S6C, respectively.

**Table S1** Percentages of VGLUT2-immunoreactive boutons in the lateral and medial parts of the LSN that were also immunoreactive for substance P (SP), tdTomato (tdTom) and/or pro-NPFF (NPFF) in two Grpr$^{CreERT2}$;Ai9 mice.

| | VGLUT2 boutons | % SP only | % tdTom only | % NPFF only | % SP and tdTom | % SP and NPFF | % SP total | % tdTom total | % NPFF total | % none |
|---|---|---|---|---|---|---|---|---|---|---|
| **Lateral** | **277.5** (248, 307) | **17.6** (14.1, 21.2) | **20.1** (19.8, 20.5) | **11.6** (10.5, 12.7) | **8.8** (8.1, 9.4) | **0.4** (0.3, 0.4) | **26.8** (22.5, 31) | **28.9** (27.9, 29.9) | **12** (10.8, 13.1) | **41.5** (35.8, 47.2) |
| **Medial** | **243.5** (173, 314) | **13.9** (11, 16.9) | **5** (2.3, 7.6) | **9.9** (12.1, 7.6) | **2.4** (2.9, 1.9) | **0** (0, 0) | **16.3** (13.9, 18.8) | **7.4** (5.2, 9.5) | **9.9** (12.1, 7.6) | **68.8** (71.7, 65.9) |
| **Pooled** | **521** (421, 621) | **15.9** (12.8, 19) | **13.3** (12.6, 14) | **10.7** (11.2, 10.1) | **5.8** (5.9, 5.6) | **0.2** (0.2, 0.2) | **21.9** (18.9, 24.8) | **19.1** (18.5, 19.6) | **10.9** (11.4, 10.3) | **54.1** (57.2, 51) |

Axons of GRPR-expressing cells are labelled with tdTomato in this mouse line, and these were much more numerous in the lateral part of the LSN. For this reason the medial and lateral parts of the LSN were analysed separately. The division between these two regions is shown in Fig S5A. Mean values from 2 mice are shown, with individual values in brackets. Column 2 shows the numbers of VGLUT2-immunoreactive boutons analysed, and columns 3-11 give the percentages for the corresponding type(s) of immunoreactivity. SP, substance P; tdTom, tdTomato; NPFF, pro-NPFF.

**Table S2** Antibodies used in this study

| Antibody | Species | Source | Catalogue # | Dilution | RRID |
|---|---|---|---|---|---|
| GFP | Chicken | Abcam | ab13970 | 1:1000 | RRID:AB_300798 |
| mCherry* | Rat | Invitrogen | M11217 | 1:1000 | RRID:AB_2536611 |
| mCherry* | Chicken | Abcam | Ab205402 | 1:5000 | RRID:AB_2722769 |
| CTB | Goat | List Biological | 703 | 1:5000** | RRID:AB_10013220 |
| NeuN-Alexa647 | Rabbit | Abcam | ab190565 | 1:1000 | RRID:AB_2732785 |
| VGLUT2 | Rabbit | Synaptic Systems | 135 408 | 1:2000 | RRID:AB_2571623 |
| VGLUT2 | Guinea pig | Millipore | AB2251 | 1:2000 | RRID:AB_1587626 |
| Calbindin | Rabbit | Swant | CB38 | 1:1000 | RRID:AB_10000340 |
| Substance P | Rat | Oxford Biotechnology | OBT06435 | 1:200 | |
| pro-NPFF | Guinea pig | M Watanabe | | 0.83 µg/ml | RRID:AB_2783015 |
| PSD-95-Alexa 647† | Alpaca | Synaptic Systems | N3702-AF647-L | 1:200 | RRID:AB_2936216 |

* The mCherry antibody also recognises tdTomato

** The CTb antibody was used at 1:200,000 for immunoperoxidase staining of brain injection sites

† The PSD-95 antibody is a camelid single-domain antibody (nanobody) conjugated to Alexa 647
